# Dietary omega-3 fatty acid deficiency from pre-pregnancy to lactation affects expression of genes involved in neurogenesis of the offspring

**DOI:** 10.1101/2022.10.14.512201

**Authors:** Vilasagaram Srinivas, Saikanth Varma, Suryam Reddy Kona, Ahamed Ibrahim, Asim K Duttaroy, Sanjay Basak

## Abstract

Maternal n-3 PUFA (omega-3) deficiency can affect brain development *in utero* and postnatally. Despite the evidence, the impacts of n-3 PUFA deficiency on the expression of neurogenesis genes in the postnatal brain remained elusive. Since postnatal brain development requires PUFAs via breast milk, we examined the fatty acid composition of breast milk and hippocampal expression of neurogenesis genes in n-3 PUFA deficient 21d mice. In addition, expression of fatty acid desaturases, elongases, signalling receptors of free fatty acids, insulin and leptin, and glucose transporters were measured. Among the genes involved in neurogenesis, the expression of brain-specific tenascin-R (TNR) was downregulated to a greater extent (∼31 folds), followed by adenosine A2A receptor (A2AAR), dopamine receptor D2 (DRD2), glial cell line-derived neurotrophic factor (GDNF) expression in the n-3 PUFA deficient hippocampus (p<0.05). Increasing dietary LA to ALA (50:1) elevated ARA to DHA ratio by ∼8 folds in the n-3 PUFA deficient breast milk, with an overall increase of total n-6/n-3 PUFAs by ∼15:1 (p<0.05) compared to n-3 PUFA sufficient (LA to ALA: 2:1) diet. The n-3 PUFA deficient brain exhibited upregulation of FADS1, FADS2, ELOVL2, ELOVL5, ELOVL6, GPR40, GPR120, LEPR, IGF1 and downregulation of GLUT1, GLUT3, and GLUT4 mRNA expression (p<0.05). Maternal n-3 PUFA deficiency affects the expression of key neurogenesis genes in the offspring with concomitant expression of desaturases and elongases genes suggesting the importance of dietary n-3 PUFA for neurodevelopment.

## 1. Introduction

For optimal growth, the fetal brain requires maternal long-chain polyunsaturated fatty acids (LCPUFAs) via placenta and breast milk *in utero* and postnatally, respectively [1-3]. Maternal PUFAs support fetal growth and brain development [4, 5]. Maternal PUFAs are important for the overall development of the fetus, including lactating mammary gland [6]. The mammary gland is an active lipid synthetic organ that secretes a large quantity of milk lipids during the 20-d lactation cycle in mice. Its development is comparable between humans and rodents. Breast milk’s fatty acids can affect the membrane’s physiochemical properties by influencing the conformation and function of the receptors, membrane-bound proteins, ion channels, and transporters in the brain [7]. The preferential postnatal deposition of LCPUFAs in the infant’s brain is mediated via breast milk. Fatty acids modulate transcription factors in controlling lipid metabolism and change the proportion of lipids in human milk [8]. The milk lipid is enriched with fatty acids, especially PUFAs. Despite low PUFA intake from the diet, breast milk enriches with an adequate amount of fatty acids indicates that the mammary gland serves as a temporary mammary pool of fatty acids in synthesizing LCPUFAs through the expression of Δ5 (FADS1) and Δ6 (FADS2) desaturases [9]. The lipid content of the diet could regulate the synthesis of PUFAs in the lactating mammary gland [10]. Data showed that a substantial regulation of mammary gland lipid synthesis occurs at gene expression levels [11]. LCPUFAs serve as precursors of second messenger signals such as eicosanoids and modulate gene expression by acting as a ligand to activate transcription factors in regulating gene expression [12, 13].

Docosahexaenoic acid, 22:6n-3 (DHA), the most predominant LCPUFA in the brain membrane, is required for fetal and neonatal brain’s structural and functional development. The signalling pathways of DHA and its metabolites are involved in neurogenesis [14]. The brain hippocampus plays a vital role in memory, and spatial navigation comprises different types of neurons. The incredible neuronal diversity in the brain results from regulated neurogenesis during its development [15]. Neurogenesis is active postnatally in the dentate gyrus of the hippocampus, whose optimal development is critical for learning and memory retention [16]. Maternal alpha-linolenic acid, 18:3n-3 (ALA), as the precursor of DHA, availability during gestation and lactation improves hippocampal development in the offspring [17]. N-3 PUFAs improve memory impairment by promoting neurogenesis [18] and their deficiency impaired neurogenesis in the rat brain [19].

Maternal n-3 PUFA deficiency decreased brain DHA content in the offspring [20, 21] and increased the percentage of total n-6 PUFAs in tissue [22]. High dietary n-6 fatty acid compromises brain DHA accretion and contributes to poor neurodevelopment [23]. DHA, obtained from the mother [1], is the key determinant for fetal neurogenesis, maturity, learning & behaviour outcome of the offspring. Maternal n-3 PUFAs are critical for the structural and functional development of the brain for effective neurotransmission [24], and neurogenesis [25-27] and others. N-3 PUFA deficiency alters neurogenesis by disrupting oligodendrocyte maturation and brain myelination [28]. The adverse n-6/n-3 PUFA ratio in gestation negatively affects brain histology [29], and hippocampal neurogenesis in mice offspring [30]. Maternal high n-6 /low n-3 fatty acids diets induce hedonic consumption in the offspring due to the programming of dopaminergic neurogenesis during embryogenesis [31]. Despite this evidence, the effects of n-3 PUFA deficiency on the expression of neurogenesis gene mediators in the postnatal brain remained elusive.

The deficiency of n-3 LCPUFA can modulate neurogenic development by affecting the brain’s gene expression in the offspring. The brain is characterized by a high gene expression level comprising 30-50% of known protein-coding genes [32]. The hippocampal expression array showed that PUFAs transcriptionally modulated genes associated with learning and appetite in mice [33]. Whether inadequate n-3 PUFA dysregulates the expression of neurogenesis genes that implicate the brain’s learning outcome is not yet known. We hypothesize that reduced n-3 PUFA delivery via breast milk could modulate the gene expression during pups’ brain development as PUFAs act as ligands for the transcription factors that control gene expression [34] including mammary glands [8, 11].

Using a mice model, we examined the effects of maternal n-3 PUFA deficiency on hippocampal expression of neurogenesis by using a PCR array in a 21d pup’s brain. In addition, we measured the impact of maternal n-3 PUFA deficiency on breast milk fatty acid composition and expression of insulin, glucose and fatty acid signalling receptors and enzymes in 21d pup’s brain.

## 2. Materials and methods

### 2.1 Diets and animals

This study was conducted after the ethical approval of the host Institute (No. NCLAS/IAEC/02/2017). The 21-d offspring brain and breast milk examined in the present investigation were obtained from our recently conducted pre-clinical trial [22]. Weanling female Swiss albino (n=30 per group) mice were kept in pairs, fed on a chow diet for a week, and maintained at 23±3°C, 55% ± 10% relative humidity with ∼12 h dark/light cycle. Post-acclimatization, mice were fed ad libitum on n-3 PUFA and a deficient isocaloric diet throughout the study. The diets were formulated according to AIN93 with modified fatty acids composition of the 10% fat source. The n-6/n-3 PUFA ratio (50:1 for n-3 deficient diet and 2:1 for n-3 sufficient diet) was achieved after blending groundnut oil, palmolein, and linseed oil (source of ALA) in varied proportions while keeping the SFA: MUFA: PUFA ratio near to ratio of 1:1.3:1 in both experimental diets [22]. Milk was collected from the dam at 9-10 PND (postnatal days). The bodyweight of the weaned mice was measured. Twenty-one days pups (n=10) were euthanized by cervical dislocation for brain (hippocampus) collection.

### 2.2 Milk and brain collection

Milking was performed as described before [35] with a few modifications. Briefly, a nursing dam (PND 9-10) with a litter of 3-4 pups was separated from pups 3h before milking, given water and the appropriate diet. Intraperitoneal injection of oxytocin (2IU/kg) was administered to the dam, followed by an anaesthetic cocktail of ketamine and xylazine of 80 µg/g and 10 µg/g of the body weight, respectively. Drops of carboxymethyl cellulose were used to keep the eyes from drying out. Mice were placed on their backs and the region around their mammary glands was cleaned with alcohol-soaked cotton. Milk was ejaculated by stroking the teat with the thumb and fingers, then squeezing the breast tissue upwards to release the milk. Milk was collected by positioning the edge of a vacuum-driven milk-collecting tube on the edge of the capillary tube, avoiding the initial few drops. This manual milk expression from teats was repeated with subsequent pairs of teats in a clockwise or anticlockwise orientation until the desired volume was achieved (200µl). For fatty acid composition, milk was kept at -80°C. Offspring were euthanized at 21-30 days to collect brains. Tissues were stored in RNA later to carry out mRNA expression.

### 2.3 Fatty acid composition of milk by gas chromatography

Milk fatty acid composition was analyzed by gas chromatography as described below. Milk (100 µl) was mixed with 5ml of methanol and chloroform (1:2) in a 15 ml screw cap glass tube and vortexed vigorously for 2 min. The mixture was filtered, collected in a fresh tube and repeatedly washed with an additional 2 ml of methanol and chloroform mixture. The whole filtrate (7 ml) was vortexed thoroughly after adding 1/5th volume of 0.22% KCl. The mixture was spun at 2000 rpm (Sigma centrifuge, Rotor ID#12159) for 2 min. The upper aqueous layer was discarded, and the lower organic phase was collected and added with C17 standard (1 mg/ml) before FAME conversion. These mixtures were incubated for 4 h at 70ºC after replacing the tube’s environment with nitrogen gas. The mixtures were cooled to room temperature, and 2 ml of H_2_O and 10 ml of petroleum ether were added. Vortexed for 2 min, the upper layer was collected into a fresh tube. This step was repeated twice under nitrogen gas. A 5 ml of NaHCO_3_ (0.2 M) was added to the above concentrate, and the upper phase was collected after vortexing for 2 min. The collected upper phase was transferred to a fresh tube containing sodium sulphate and stored at -20ºC for analysis. The composition of fatty acids was analyzed by gas chromatography (Perkin Elmer Clarus 680) employing a flame ionization detector with an SP2300 fused silica capillary column (30 m × 0.25 mm × 0.2 μm, Supelco, USA). The standards with recognized retention times were used to identify the samples (Nuchek, USA).

### 2.4 Neurogenesis pathway analysis by qRT-PCR array

RT^2^ profiler PCR arrays combine real-time PCR (sensitivity, reliability) and microarray (multigene profiling) to simultaneously detect the expression of several genes of a targeted pathway. Genes involved in the mouse neurogenesis pathway were studied by RT^2^ profiler PCR array (**Supplementary Table 1**). Briefly, total RNA was extracted from the hippocampus of the brain, followed by gDNA elimination and cDNA conversion as per manufacturers’ instruction (Qiagen, Cat# 74104 and Cat# 330401). A mouse neurogenesis PCR array kit (#PAMM-404AZA, Qiagen) was employed to measure the mRNA expression of genes involved in this pathway using real-time PCR (ABI 7500). The changes in the mRNA fold expression difference were calculated by the ΔΔCt method, where (2^−ΔΔ^^Ct^) is the normalized mRNA expression of (2^−ΔCt^) of n-3 deficient over normalized mRNA expression (2−^ΔCt^) of the n-3 sufficient group. The minimum cut-off expression difference was considered ≥ 2 folds.

**Table 1:**
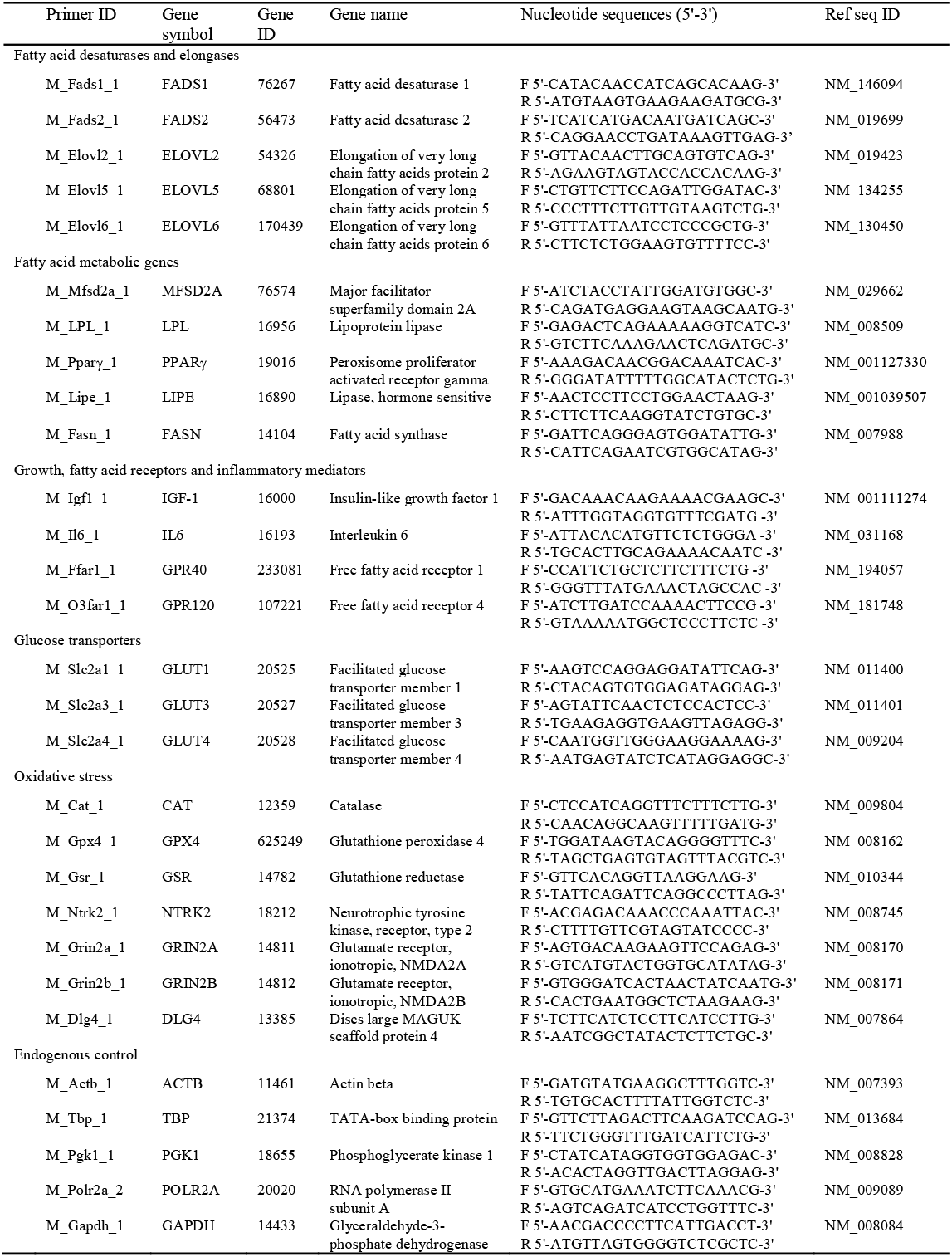
Gene pathways and predesigned SYBR green I primers used for the mRNA expression analyses of mice brain

### 2.5 Gene expression analyses by real-time PCR

Brain tissue (50mg) was finely minced and transferred into 1 ml of cold Tri reagent (# T9424, Sigma) containing 2 mm zirconia beads (#11079124Zzx, Biospec). Tissues were homogenized for 1-2 min (Mini Bead beater, Biospec) and spun at 12,000g for 10 min to eliminate cell remnants (4ºC). Total RNA was purified as per supplier instructions. Isolated total RNA underwent gDNA elimination (#AMPD1, Sigma) and quality and quantity evaluation by spectrophotometer (#ND1000, Thermo Scientific). Total RNA (1µg) was converted to cDNA (#1708891 Bio-Rad), and relative mRNA expression was measured using pre-designed primers compatible with SYBR green I chemistry (gene ID, pathways and nucleotide sequences are mentioned in **Table1)**. The tissue-specific, accurate endogenous control was applied in mRNA expression analysis of genes by averaging the expression of multiple control genes using Norm Finder software (v20). The expression of mRNA was calculated after obtaining Ct values using the comparative delta Ct method (2−^ΔΔCt^) after normalized with the most stable endogenous control. Data were presented as fold change over control after normalizing with the endogenous control gene.

### 2.6 DNA methylation analyses

Global DNA methylation was estimated in the hippocampus. The genomic DNA was purified from the hippocampal region of the brain tissues (∼10-12mg) using the smart extract-DNA extraction kit as per the supplier’s instruction (Cat# SKDNEX-100, Eurogentec, Belgium). The quantity of total genomic DNAs (gDNA) was adjusted equally across the samples. Input gDNA (100ng) was used as a template to calculate methylation percentage by Methyl flash kit (Cat# P-1030, Epigentek, USA) as described previously [22].

### 2.7 Statistical analyses

Unpaired Student’s t-test was employed to compare two groups using GraphPad Prism v.8. Statistical significance was considered when the p-value was less than 0.05. The assays were carried out independently repeated times, as indicated in the figure legends. The values are expressed as a mean ± standard error of the mean (SEM).

## 3. Results

### 3.1 The maternal n-3 PUFA deficiency changes brain weight in the pups

As reported earlier [22], the intake of an n-3 PUFA deficient diet did not influence a change in pre-pregnancy bodyweight in the dam. The cumulative food intake (Kcal) and feed efficiency (%) did not differ over a period of nine-weeks in dams (data not presented). The body weight and liver weights showed no significant difference in 21-d pups (**Table 2**). The n-3 PUFA deficient brain weight was increased marginally in these pups (p<0.05). However, brain to bodyweight ratio was similar between the two groups (**Table 2**).

**Table 2:** Body weight and organ weight of 21d pups ^#^

| Weight (g) | n-3 PUFA sufficient | n-3 PUFA deficient | Mice no. | p value |
| --- | --- | --- | --- | --- |
| Body weight | 6.956 ± 0.352 | 7.249 ± 0.348 | n=18 | 0.561 |
| Liver | 0.28 ± 0.014 | 0.278 ± 0.025 | n=18 | 0.952 |
| Brain | 0.357 ± 0.004 | 0.370 ± 0.004 | n=18 | 0.046* |
| Brain/body weight ratio | 0.051: 1 | 0.051: 1 | n=18 | - |

<sup>#</sup> Values (in grams) are expressed as mean ± SEM. n= number of mice from each group.
Paired t-test between n-3 deficient and n-3 sufficient diet group; \* p<0.05;

### 3.2 The maternal n-3 PUFA deficiency affects the expression of the neurogenesis genes in the offspring’s brain

We investigated the expression of neurogenesis gene mediators in the offspring brains by a qRT-PCR array. **Table 3** summarizes the fold expression changes of the genes associated with mouse neurogenesis pathway in n-3 PUFA deficient hippocampus over the n-3 PUFA sufficient mice. The altered genes (up or down) are adjusted to the expression cut-off > 2.0 folds as presented between these two groups (**Fig.1)**. Seven key genes such as TNR, A2AAR, DRD2, GDNF, TGFB1, TH, and POU3F3 were significantly downregulated in the n-3 PUFA deficient brain. Among these genes, the most notable change showed a substantial decrease in TNR mRNA levels by ∼31 folds, a brain-specific gene expressed in the nervous system. The brain-specific expression of genes such as A2AAR, DRD2, and GDNF was decreased by ∼3.03, ∼2.33, and ∼2.51 folds, respectively, in the deficient hippocampus (**Fig.1)**. These genes are associated with differentiation, adhesion, migration, survival, neurotransmitter synthesis, release, and signalling of neuronal cells present in the brain. Contrary to the downregulation, the expression of eight mRNA, such as BMP4, EFNB1, EGF, NEUROG2, NOG, PAX5, SLIT2, and SOX2 was upregulated in n-3 PUFA deficient offspring brain. Among these, a brain-specific NEUROG2 expression was activated in adult hippocampal neurogenesis.

**Table 3:**
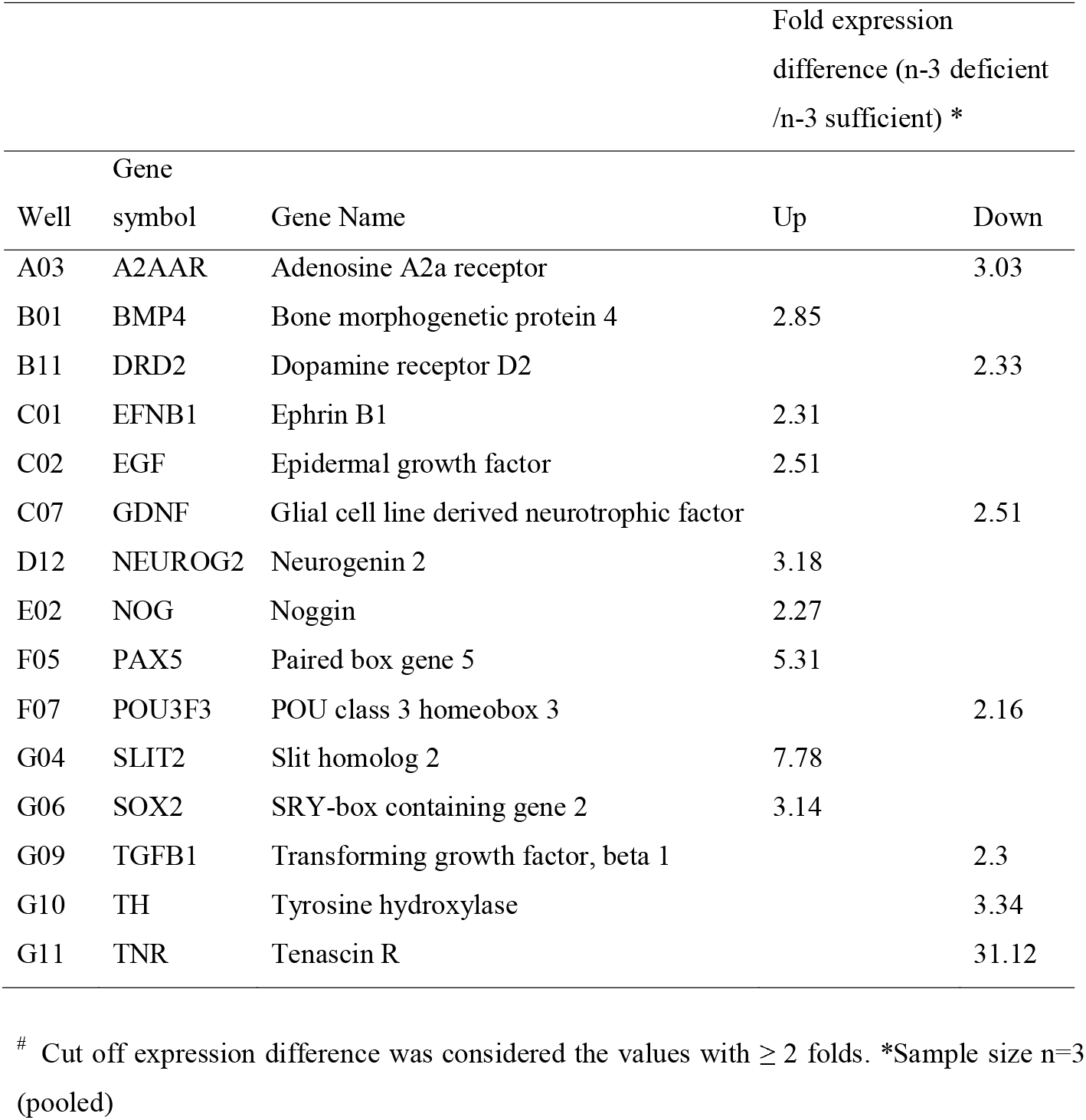
Effect of maternal n-3 PUFA deficiency on neurogenesis gene expression pathway in pup’s brain tissue^#^

**Fig 1.**
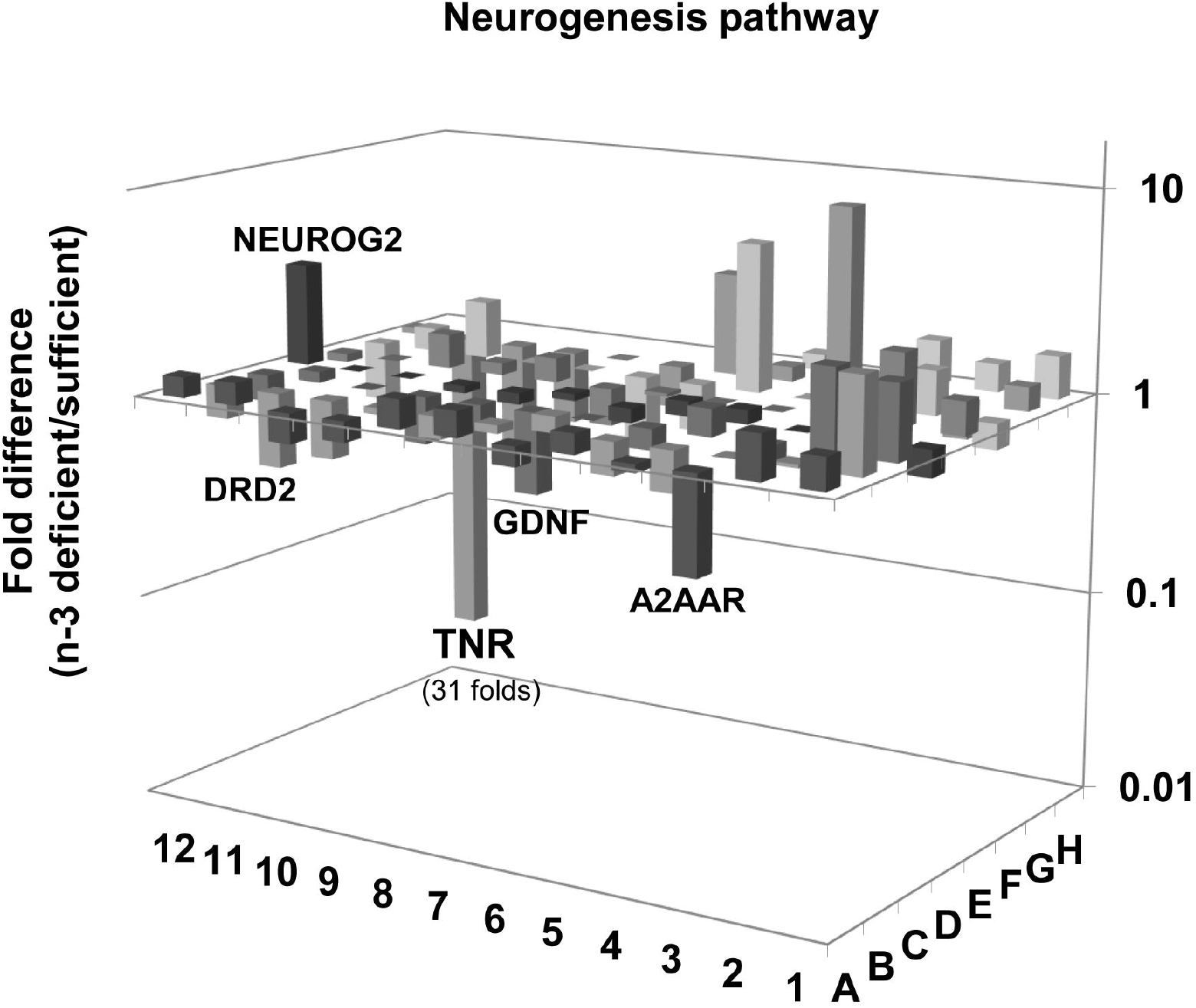
Effect of maternal n-3 PUFA deficiency on neurogenesis pathway analysis in the hippocampal brain of 21-d offspring [n=3 (pooled)/group]. Detailed experimental condition is explained in the methodology section. Each 3-D bar shows mRNA fold expression either up or down-regulation caused due to n-3 PUFA deficiency after normalizing with housekeeping genes.

### 3.3 Endogenous control gene of the brain was selected by determining the stability value of candidate gene expression

Norm Finder was employed to measure the expression stability of the five potential endogenous control genes such as ACTB, TBP, PGK1, POLR2A, and GAPDH in the hippocampal brain of 21-d male Swiss albino mice (n=5) after quantification of their mRNA expression by qRT-PCR. Log dCt of the respective genes was used to assess the expression stability using Norm Finder software (June 2014). The analysis revealed that the five genes exhibited a stability value ranging between 0.491 to 0.901 (**Fig.2**), where ACTB has the lowest stability value by Norm Finder. A lower stability value indicates higher consistency in expressing Ct across the replicates when used as an endogenous control gene in hippocampal brain tissue.

**Fig 2.**
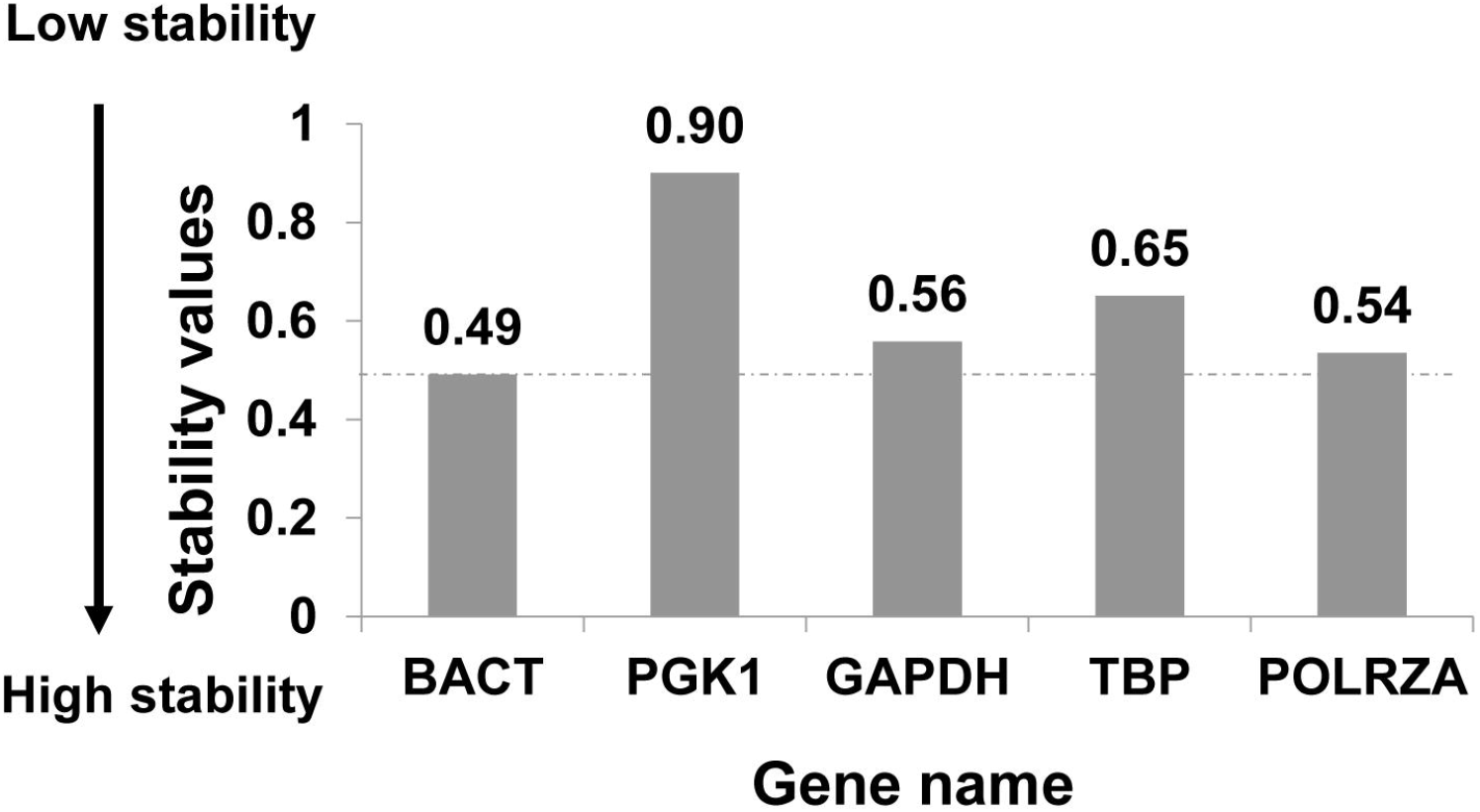
The stability value of mRNA expression of endogenous control genes in the hippocampal brain of 21-d male Swiss albino mice (n=5/group). As mentioned in the text, the mRNA expression of genes was quantified by qRT-PCR and analyzed by Norm Finder.

### 3.4 Brain fatty acid desaturase and elongases expression was upregulated in offspring due to maternal n-3 PUFA deficiency

A reduced supply of maternal PUFAs could affect the expression of genes encoding for the enzymes responsible for the endogenous conversion of PUFAs into LCPUFAs. Therefore, we investigated the mRNA expression of genes involved in fatty acid desaturation and elongation in the offspring hippocampal tissue. Desaturation and elongation activities were increased in the n-3 PUFA deficient brain via a significant (p<0.05) upregulated expression of FADS1 (∼2.05), FADS2 (∼1.58), ELOVL2 (∼1.55), ELOVL5 (∼1.63), and ELOVL6 (∼1.73) mRNAs **(Fig.3 A-B)**. Thus, the maternal deficiency of n-3 PUFA marked a significant alteration in the expression of fatty acid desaturases and elongase genes in the offspring’s brain.

**Fig 3.**
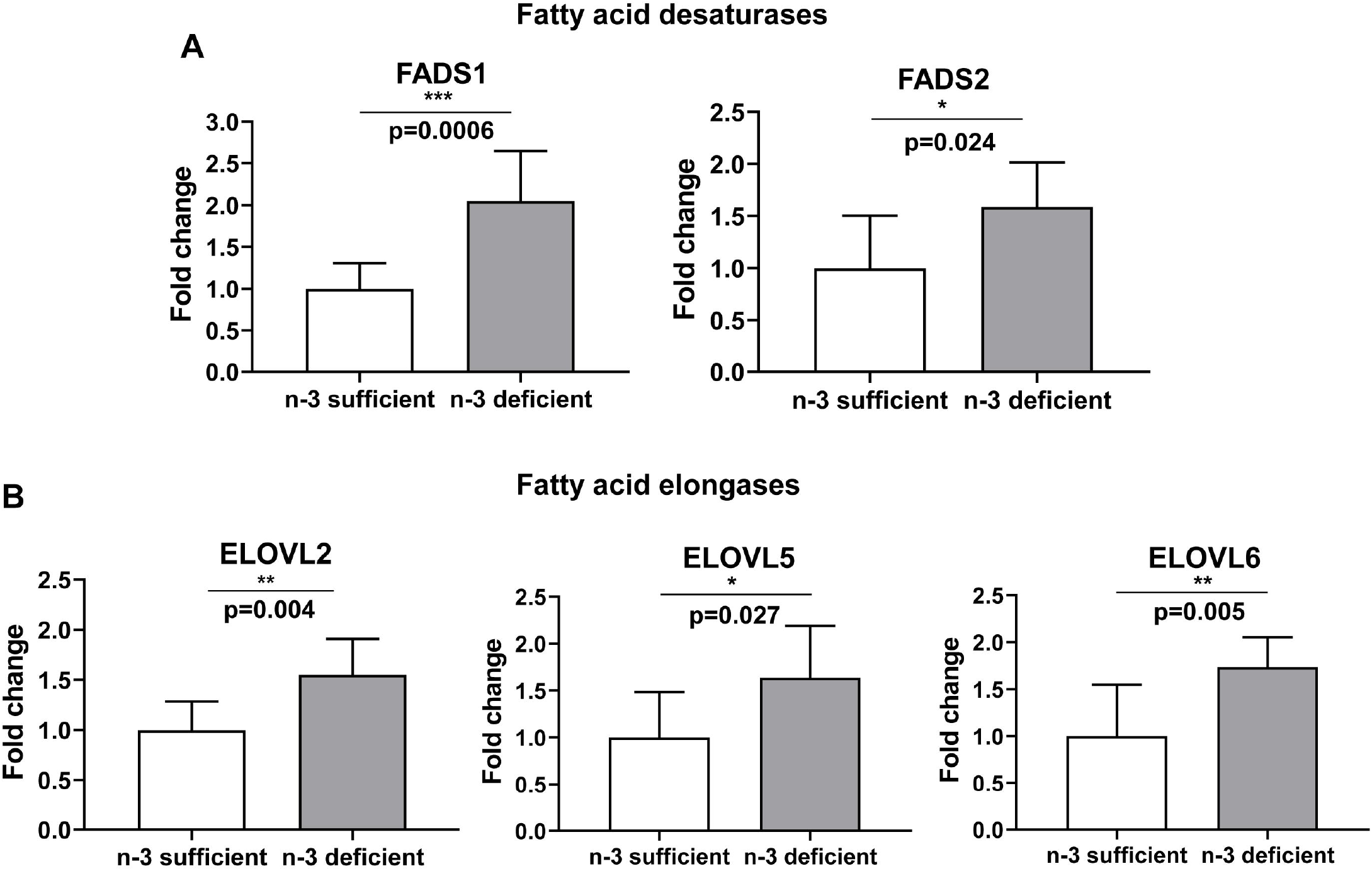
Effect of maternal n-3 PUFA deficiency on mRNA expression of fatty acid chain desaturation and elongation genes in the pup’s hippocampal brain. (A) Fatty acid desaturases, FADS1 and FADS2 (B) Fatty acid elongases, ELOVL2, ELOVL5, and ELOVL6 expression were measured in the hippocampus of 21d pups born to n-3 PUFA sufficient and deficient fed dams. The mRNA expression of gene expression was measured using RT-qPCR, normalized with the mRNA expression of ACTB, and expressed as fold changes over n-3 PUFA sufficient groups (n=8/group). Individual p-values are indexed in figures. *p<0.05 vs n-3 sufficient (Student’s t-test).

### 3.5 Brain insulin & leptin signalling, fatty acid signalling mediators’ expression were upregulated in the offspring due to pre-pregnancy n-3 PUFA deficiency

A deficient supply of maternal n-3 PUFAs could affect the expression of fatty acid signalling sensors in the brain tissue. Therefore, we investigated the mRNA expression of genes involved in lipoprotein transporters, fatty acid signalling receptors, and pro-inflammatory markers in 21-d offspring hippocampal tissue. The n-3 PUFA deficient brain showed a significant increase (p<0.05) in the mRNA fold expression of GPR40 (∼2.9), GPR120 (∼2.8), LEPR (∼2.8) and IGF1 (∼2.2), respectively (**Fig.4 A-B)**. However, these two groups’ expression of pro-inflammatory IL6, lipoprotein transporters MFSD2A, PPARγ and LPL were comparable (data not shown). Thus, the gestational deficiency of n-3 PUFA marked a significant alteration in the brain’s fatty acid signalling and receptor expression in the young pups.

**Fig 4.**
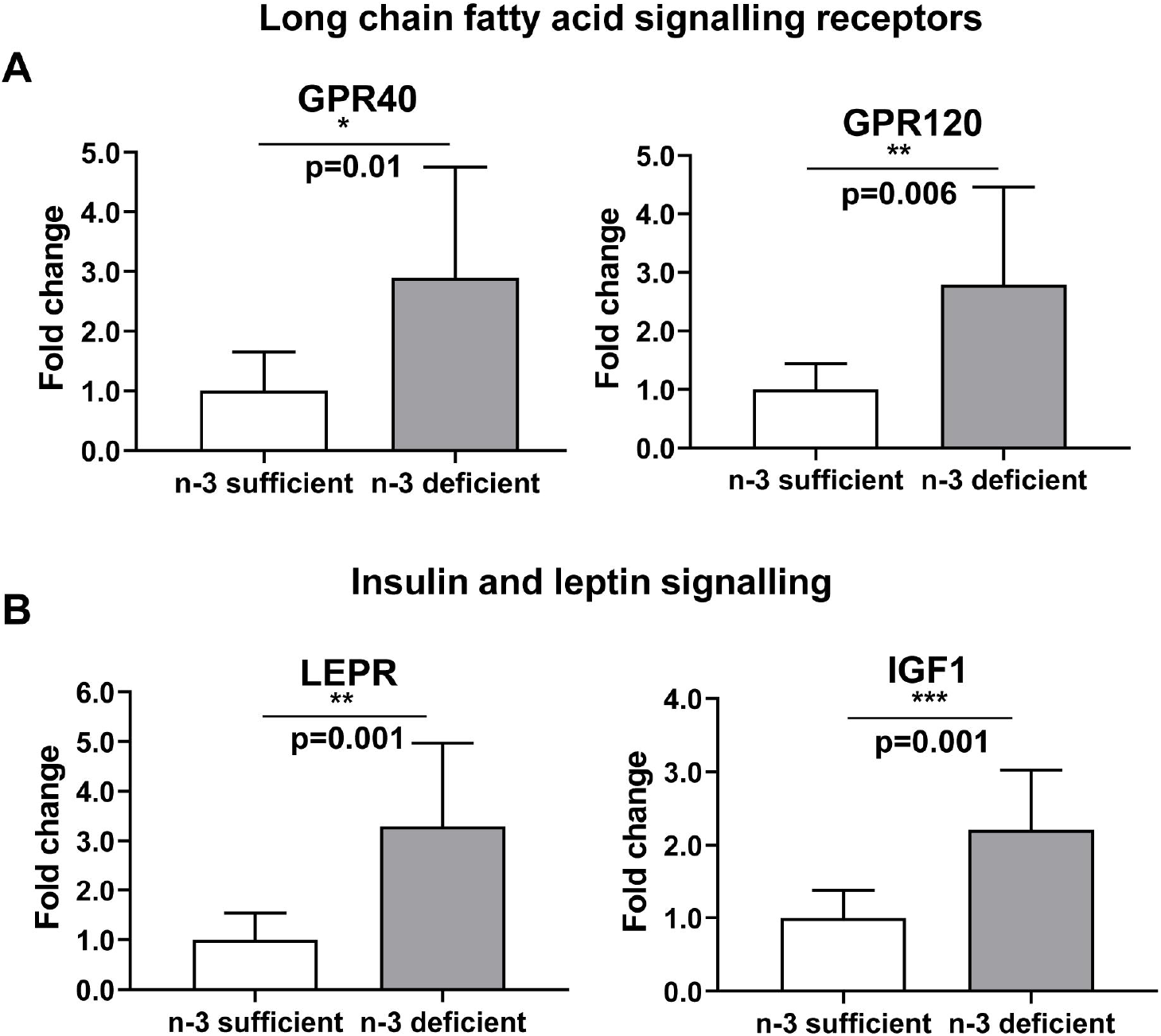
Effect of maternal n-3 PUFA deficiency on the mRNA expression of genes, (A) long chain fatty acid signalling receptors, (B) insulin and leptin signalling mediators in the pup’s hippocampal brain. The mRNA expression was measured using RT-qPCR, normalized with ACTB, and expressed as fold changes over n-3 PUFA sufficient groups (n=9/group). Individual p-values are indexed in figures. *p<0.05 vs n-3 sufficient (Student’s t-test).

### 3.6 Brain glucose transporters expression was downregulated in the offspring due to maternal n-3 PUFA deficiency

The primary energy needs of the brain are met by glucose via glucose transporters (GLUTs). Therefore, we investigated the expression of major GLUTs expressed in the brain. The fold changes in the mRNA transcripts of GLUT1 (∼2.39), GLUT3 (∼2.96), and GLUT4 (∼2.07) were considerably down-regulated in the brains of n-3 PUFA deficient pups (p<0.05, **Fig.5)**.

**Fig 5.**
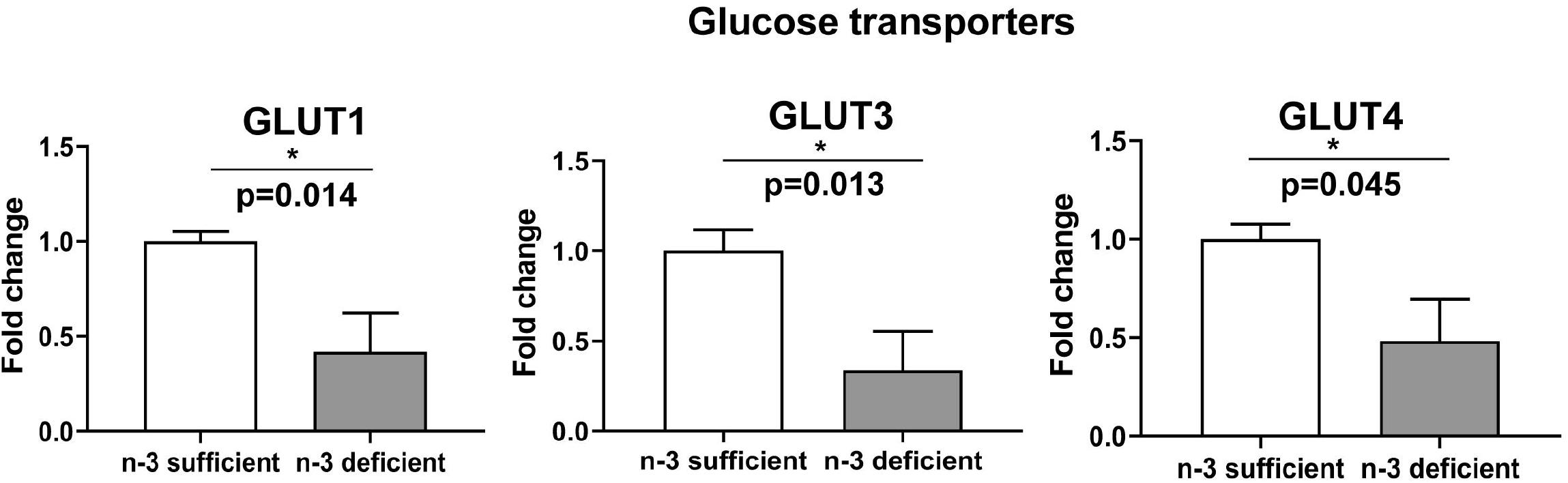
Effect of n-3 PUFA deficiency on glucose transporters (GLUTs) in the hippocampus of 21d pups. The gene expression was measured in brain tissue of pups born to n-3 PUFA sufficient and deficient fed dams. The mRNA expression was analyzed using quantitative real-time qRT-PCR normalized to ACTB as endogenous control. Fold change of gene expression was calculated according to the ^2-ΔΔ^ Ct as indicated in the method. Data represented in mean± SEM (n=6/group) *p-values <0.05 vs. n-3 sufficient.

### 3.7 Epigenetic DNA methylation and expression of oxidative stress biomarkers were unaffected in the pups’ brain

Nutritional deficiency of n-3 PUFA during gestation and lactation may induce epigenetic changes in neuronal cells during fetal brain development. The epigenetic changes were measured by scoring the global DNA methylation percentage, as reported before [22]. The maternal n-3 PUFA deficiency did not affect the levels of 5-methylcytosine (%) in the offspring hippocampus (n-3 PUFA suff. vs def.: 31.21 ± 6.61 vs 32.65 ± 7.72, p>0.05). Since n-3 fatty acids are reported for their antioxidant activity, their protective functions were measured by mRNA expression of CAT, GPX4, GSR, NTRK2, GRIN2A, GRIN2B, and DLG4 genes. The maternal n-3 PUFA deficiency did not affect the expression of genes associated with antioxidant activity in the hippocampus of the offspring (data not shown).

### 3.8 Breast milk fatty acid composition was affected due to maternal n-3 PUFA deficiency

Maternal PUFAs are delivered to the fetus via the breast milk of the mother. Therefore, breast milk fatty acid composition was evaluated to trace the probable estimate of PUFAs delivery to the fetal brain. The total fatty acid composition (% nmole) of breast milk obtained on the 10^th^ day from lactating dams is presented in **Table 4**. Maternal n-3 deficiency caused a significant decrease in n-3 LCPUFAs content of DHA (C22:6 n-3) and DPA (C22:5 n-3) in the breast milk by ∼6.0 (p=0.0004) and ∼6.5 folds (p<0.0001) respectively as compared to n-3 sufficient dam. This was accompanied by a progressive increase in the proportion of n-6 LCPUFAs levels such as ARA (C20:4 n-6) and ADA (C22:4 n-6) by ∼1.2 (p=0.079), and ∼2.0 (p=0.0004) folds respectively (**Table 4**).

**Table 4:** Breast milk fatty acid composition (% nmole) at 10^th^ day lactation of dams fed with n-3 PUFA sufficient and deficient diet

| Fatty acids<br>(nmol %) | n-3 sufficient (n=7) | | n-3 deficient (n=7) | | p value | Fold<br>change<sup>\$</sup> |
| --- | --- | --- | --- | --- | --- | --- |
|  | Mean | SEM | Mean | SEM |  |  |
| C12:0 | 12.25 | 0.59 | 14.61 | 0.71 | 0.025* | - |
| C14:0 | 13.73 | 0.46 | 15.86 | 0.55 | 0.011* | - |
| C14:1 | 0.58 | 0.03 | 0.52 | 0.03 | 0.208 | - |
| C16:0 | 24.22 | 1.29 | 22.89 | 0.63 | 0.375 | - |
| C16:1 | 4.38 | 0.28 | 1.71 | 0.74 | 0.005* | - |
| C18:0 | 3.54 | 0.13 | 3.07 | 0.12 | 0.025* | - |
| C18:1 | 24.08 | 0.61 | 25.80 | 0.96 | 0.156 | - |
| C18:2 n-6 | 10.91 | 0.27 | 13.32 | 0.81 | 0.016* | - |
| C20:4 n-6 | 0.71 | 0.05 | 0.89 | 0.07 | 0.079 | 1.2 (+) |
| C22:4 n-6 | 0.14 | 0.01 | 0.28 | 0.02 | 0.0004*** | 2.0 (+) |
| C18:3 n-3 | 3.91 | 0.18 | 0.81 | 0.45 | <0.0001*** | 4.8 (-) |
| C20:5 n-3 | 0.62 | 0.03 | ND | ND | - | - |
| C22:5 n-3 | 0.46 | 0.01 | 0.07 | 0.05 | <0.0001*** | 6.5 (-) |
| C22:6 n-3 | 0.39 | 0.04 | 0.06 | 0.04 | 0.0004*** | 6.0 (-) |
| <sup>1</sup>ΣLC n-6 PUFA | 0.85 | 0.06 | 1.17 | 0.09 | 0.204 | 1.3 (+) |
| <sup>2</sup>ΣLC n-3 PUFA | 1.47 | 0.08 | 0.13 | 0.09 | <0.0001*** | 11.3 (-) |
| ARA: DHA | 1.8:1 | - | 14.8:1 | - | - | 8.2 (+) |
| ΣLC(n-6): (n-3) | 0.58 | - | 9.0 | - | - | 15.5 (+) |
| Σ(n-6): (n-3) | 2.18 | - | 15.4 | - | - | 7.06 (+) |

<sup>\$</sup> Fold change in fatty acid concentration are depicted by increase (+) or decrease (-) over n-3 sufficient group.
ND = non-detectable
<sup>1</sup> ΣLC n-6 PUFA is the sum of C20:4 n-6 and C22:4 n-6
<sup>2</sup> ΣLC n-3 PUFA is the sum of C20:5 n-3, C22:5 n-3 and C22:6 n-3
Paired test between n-3 deficient and n-3 sufficient group; Mean ± SEM, \* p<0.05, \*\*\* p<0.0005;

The n-6/n-3 PUFA ratio of 2:1 in the maternal diet of the n-3 PUFA sufficient dam was reflected around 2.8:1 in the breast milk, indicating a closely linear dietary conversion of n-3 PUFAs into the breast milk n-3 LCPUFAs (18:2n-6/18:3n-3=10.91/3.91). In n-3 PUFA deficient dam, the dietary n-6/n-3 PUFA (50:1) ratio was correlated to ∼16:1 in the breast milk (18:2n-6/18:3n-3=13.32/0.81), indicates a still increased levels of n-6 PUFAs in the n-3 deficient dam’s breast milk. Thus, a proportion of n-3 PUFAs supplied by the maternal diet to breast milk was efficiently converted in n-3 PUFA sufficient dam but not in the n-3 deficient mice.

The increasing presence of LA to ALA (50:1) in the diet significantly elevated ARA to DHA ratio by ∼8 folds in the n-3 PUFA deficient breast milk (**Table 4**). The ratio of ARA to DHA was found ∼1.8:1 in n-3 PUFA sufficient and 14.8:1 in n-3 PUFA deficient breast milk. Moreover, the endogenous conversion of n-3 LCPUFAs from dietary ALA (C18:3 n-3) was decreased by ∼2.4 folds in n-3 deficient breast milk (n-3 LCPUFA/18:3 n-3 PUFA:1.47/3.91= 0.38/1 n-3 suf. vs. 0.13/0.81 = 0.16/1 n-3 def.), whereas the conversion of n-6 LCPUFAs from dietary LA (C18:2 n-6) was found closely similar in between n-3 PUFA sufficient and deficient dams (n-6 LCPUFA/18:2 n-6: 0.85/10.91 = 0.07/1 n-3 suf. vs. 1.17/13.32 = 0.08/1 n-3 def.). Compared to n-3 PUFA sufficient dam, the conversion efficiency to form n-3 LCPUFAs from their dietary precursors remained lower with a significant surge in ARA to DHA ratio in n-3 PUFA deficient breast milk.

## Discussion

This is the first study that demonstrated a maternal n-3 PUFA deficient diet from preconception until lactation affected the expression of neurogenesis genes in the hippocampal brain of the 21-d offspring mice, particular with a significantly reduced expression of Tenascin R (TNR) gene. Moreover, increased expression of genes that encodes fatty acid desaturases, elongases, insulin growth factor-1, leptin receptor, free-fatty acid receptor, and decreased expression of glucose transporters was also observed in the hippocampus of the offspring born to n-3 PUFA deficient dam. Breast milk fatty acid composition reveals that n-3 PUFA deficiency not only lowered the delivery of n-3 LCPUFAs to the fetus but also decreased the efficiency of endogenous conversion of n-3 LCPUFAs from their precursors. Collectively, these data suggest that maternal n-3 PUFA deficiency affects hippocampal neurogenesis in mice by dysregulated gene expression in the post-weaned offspring due to inadequate mammary supply of LCPUFAs.

The mechanism that involved a drastic reduction of TNR expression in the n-3 PUFA deficient mice brain (**Fig.1**) is not known yet. TNR is an extracellular matrix (ECM) glycoprotein exclusively present in the nervous system. Even though TNR expression decreases postnatally in oligodendrocytes, its expression continues in the mature neuron of the rodent [36]. Decreased TNR expression may affect migration, adhesion and differentiation of neuronal cells associated with its development. Recent data suggest that n-3 PUFA deficiency alters brain function by disrupting oligodendrocytes maturation and brain myelination during neurodevelopment [28]. Since TNR acts as an autocrine factor for oligodendrocytes differentiation, a reduced expression of TNR could affect this differentiation [37], which was corrected by the addition of TNR [38]. TNR plays a role in neuroprotection by modulating microglial function [39] and its expression was inhibited by TNF-α produced by activated microglia [40]. The TNR deficient mice showed learning and memory deficit, reduced exploratory behaviour, impaired fine motor function [41], and increased anxiety due to hippocampal excitability [42]. TNR is a critical component in the brain’s ECM and appears to maintain optical distance between neuronal cells. The TNR deficient mice showed a reduced density of perineuronal nets and altered synaptic activity [43]. Thus, a drastic decrease in TNR expression due to n-3 PUFA deficiency could alter the neuronal density and microglial inflammation in the offspring brain and may increase susceptibility to neuroinflammation induced by spatial memory impairment [44].

Adenosine A2A receptor (A2A), a class of G-protein coupled receptors in the brain, is involved in neurotransmission that controls motor skills [45]. Neuronal overexpression of A2A is neuroprotective [46] indicating a reduced expression in the n-3 PUFA deficient brain (**Fig.1**) devoid of such protection. DHA chain enhanced G-protein activation by A2A [47]. A2A is located inside the lipid bilayer of the acyl chain derived from DHA and ARA. Thus, maternal n-3 PUFA deficiency may influence the motor skills of the offspring due to inadequate DHA supply (**Table 4**) and reduced A2A expression. The n-3 PUFA deficiency selectively impaired motivation in mice by inhibiting DRD2 (dopamine receptor D2)-expressing neurons [48]. Inadequate levels of n-3 LCPUFAs in breast milk and reduced expression of DRD2 in the brain observed in our study indicate a defect in the dopaminergic system of the n-3 PUFA deficient brain that may increase offspring’s risk of developing impulse-control and attention deficit behaviour. A significantly downregulated expression of GDNF and reduced breast milk DHA was observed in the n-3 PUFA deficient mice. A simultaneous increase in DHA content and GDNF levels in the hippocampus of DHA-supplemented rats [49] suggest that n-3 PUFA deficiency could affect neurogenesis by modulating GDNF expression in the hippocampal brain of the mice. Maternal n-3 PUFA deficiency also overexpressed genes such as BMP4, EFNB1, EGF, NEUROG2, NOG, PAX5, SLIT2, and SOX2 in the hippocampal brain of the offspring. Among these genes, activated neurogenin expression specifies the hippocampal role in adult neurogenesis [50] that might have a compensatory response to maintain homeostasis of brain functionalities. The expression of remaining genes encodes transcription factors and growth factors that control multiple functions in diverse tissues, including the brain.

Deficient n-3 PUFA-mediated suppression of TNR expression may affect differentiation in brain cell maturation and delay neuronal maturity. A reduced perineuronal density due to lowered TNR expression could impede synaptic connectivity and prevent further adding domains for improving learning and memory function. Brain tissue is characterized by high levels of gene expression comprising 30-50% of known protein-coding genes [32]. The rodent TNR gene exhibited 93% homology of the protein-coding region with human TNR [51], indicating a reduced expression due to maternal n-3 PUFA deficiency could be implicated in the neurogenesis of humans.

A significant lower concentration of ALA and/or DHA in the breast milk could impair brain energetics associated learning behaviour of the n-3 PUFA deficient offspring. The brain can metabolize fat as energy since neonates are nourished with lipid-enriched breast milk during the early stage of its development [24]. ALA and LA have multiple roles in the brain, including as energy substrates for peroxisomal beta-oxidation for mitochondrial fuel and as substrates for endogenous lipid synthesis within a blood-brain barrier. LA and ALA constitute structural lipids in normal brain function, ranging around 1% LA and 0.2% ALA in the balanced proportion. The uptake of LA and ALA and its utilization in the brain may be affected by the proportion of LA and ALA present in the diet and breast milk, as evidenced in our study. Rats fed with a high LA diet over two generations had significantly lowered brain ATP concentration than an ALA-rich diet implicated an altered learning outcome [52]. The maternal n-3 PUFA deficiency significantly reduced the expression of glucose transporters such as GLUT1, GLUT3 and GLUT4 in the offspring hippocampus (**Fig.5**). The changes in brain energy homeostasis induced by n-3 fatty acid deficiency could result from functional alteration of glucose transporters [53]. An increase in GLUT3 immunoreactivity promotes adult neurogenesis in the dentate gyrus of the hippocampus [54]. Thus, the n-3 PUFA deficiency can impair neurotransmission partly by suboptimal brain energy metabolism, i.e., glucose entry into the brain [24]. Despite comparable calorie intake and feed efficiency, maternal n-3 PUFA deficiency significantly altered insulin and leptin signalling by increased expression of LEPR and IGF1 in the offspring brain (**Fig.4B**). The increased n-6/n-3 fatty acids, ARA to DHA ratio in breast milk, IGF1 levels observed in the brain of the present study indicate offspring’s susceptibility to altered body fat development [55]. The increased n-6/n-3 fatty acid ratio significantly alters the peripheral marker of energy homeostasis by modulating leptin [56]. Leptin affects postnatal brain development by modulating neurogenesis, axon growth, and synaptogenesis [57]. Leptin promotes hippocampal neurogenesis in vivo and in vitro [58]. Increased LEPR expression due to n-3 PUFA deficiency could affect hippocampal neurogenesis in the offspring due to the lower availability of leptin.

ARA and DHA ratio in breast milk could have skewed for multiple reasons, including a high n-6/n-3 PUFA diet, endogenous conversion of PUFAs and increased expression of desaturases and elongases, as observed in our study. LCPUFAs accumulate in the fetal brain during the last third of pregnancy at a rate of ∼ 67mg/day. DHA and ARA make up roughly 20% and 15% of the brain’s fatty acids, respectively. Approximately 1.5 years after birth, the brain’s total volume surges to 80% of its maximal capacity [59], close to the 21-d mice pups. Maternal plasma & placenta [22], and breast milk fatty acid composition observed in this study are closely reflected in maternal n-6 and n-3 PUFA ratio in the diet. Data suggest that early infant adiposity is positively associated with the n-6 to an n-3 fatty acid ratio in human milk independent of BMI [60]. The decreasing levels of LCPUFAs (ARA and DHA) in breast milk correlate with pregnant women with obesity [61]. The offspring derived from n-3 PUFA deficient dam predispose the adult risk of gestational obesity. Thus, the n-6/n-3 PUFA ratio of breast milk correlates with maternal diet before pregnancy which could be a key predictor of later health and wellbeing of an individual. However, both DHA and ARA are involved in early brain development [62]. Since the metabolism of ARA is affected by n-3 PUFA deficiency and thus this may also affect brain development

Lactating women who ingest ALA-rich-chia seeds oil increased the DHA content of breast milk [63]. *In vivo* data showed that free (unesterified) ALA and DHA can traverse the blood-brain barrier from the circulatory plasma lipid pool. The endogenous conversion of ALA to DHA can occur inside the central nervous system from the supply of an ALA-rich diet. The endogenous fatty acid stores influence breast milk fatty acid composition rather than immediate intake [64]. In this way, sufficient DHA is ensured for the brain from an ALA-rich diet, which is comparable to DHA-fed rats [65]. The endogenous fatty acid stores are extremely important to start with pregnancy as majorities of the milk lipids (∼60%) are derived from the maternal store [66]. Thus, maternal n-3 PUFA deficiency risks delivering lower n-3 LCPUFAs to the fetus and decreases the efficiency of endogenous conversion of n-3 LCPUFAs from their precursors (**Table 4**).

Typically, ARA to DHA ratio remained relatively constant (1.5-2.0 to 1.0) in human breast milk globally [67]. The maternal n-3 fatty acid deficiency deranged the normal ARA: DHA ratio (2:1) of the breast milk observed in our study, which can alter the growth trajectory of the developing pups [61]. The presence of ARA in a higher amount may support growth-promoting activities and the production of eicosanoids. The structure-function and metabolism of the brain depend on the optimal levels of ARA and DHA and the interplay of their metabolites [68]. The recycling (de-esterification–re-esterification) of DHA and ARA in the brain is independently mediated by DHA- and ARA-selective enzymes. The ARA-dependent processes can be altered differentially than DHA-mediated processes during dietary deficiency of n-3 PUFA, as evidenced in our study. Thus, increased ARA metabolism compensated for DHA deficiency in the n-3 PUFA deficient breast milk. However, more data are required to understand the impact of the n-6/n-3 PUFA ratio on the regulation of DHA-selective iPLA2 and COX-1 or ARA-selective cPLA2/sPLA2 and COX-2 and their effects on brain function and neuroinflammation [62]. Reduced brain n-3 PUFAs change the membrane fluidity and alters nerve communication due to an imbalance between less pro-inflammatory n-3 PUFAs and more pro-inflammatory n-6 PUFAs derived eicosanoids and may alter the signalling actions via GPR120 receptors [69]. The maternal n-3 PUFA deficiency significantly upregulated the expression of GPR40 and GPR120 in the offspring’s brain (**Fig.4A**). It might promote altered signalling in the energy homeostasis and inflammation [69] in n-3 PUFA deficiency.

The increased expression of FADS1 and FADS2 in the brain might attempt to compensate for the reduced supply of DHA to the brain via breast milk, as evidenced by a high n-6/n-3 PUFA diet previously [70]. Increased expression of desaturases and elongases in the present study was similarly observed in rat liver (not brain) fed with n-3 PUFA deprived diet [71]. The tissue-specific discrepancies in the expression of these PUFA conversion enzymes could be due to the composition of the n-3 PUFA deficient diets (0.2 % vs 0.13% ALA), stage in brain maturity, and exposure window between these studies.

We extensively measured gene expression (**Sup. Table 1**) in brain tissue by qRT-PCR. The tool received huge attention in the recent pandemic due to its diagnostic application. Even though the technique is sensitive and powerful for measuring gene expression, the expression of reference genes (endogenous control gene or housekeeping gene) for each tissue contributes a powerful role in determining the mRNA expression of genes accurately [72]. Normalizing the test genes with the most stable endogenous control in mice brains (**Fig.2**) might have reflected the closest changes in the gene expression with humans due to genome similarity between these two species.

## Conclusions

The study has shortcomings, including the lack of mammary gland fatty acid compositional data and the protein expression of key genes in the hippocampus. All of these occur due to limited tissue availability in mice. Whether the expression of neurogenesis genes was affected due to n-3 PUFA deficiency in pregnancy or lactation could not be ascertained since the deficiency was continued throughout the study. Nevertheless, present data suggest for the first time that downregulated expression of TNR, A2AAR, DRD2, and GDNF, along with reduced DHA levels in the breast milk, could affect postnatal neurogenesis in the n-3 PUFA deficient offspring brain. The n-3 PUFA deficiency further deranged the breast milk’s normal ARA: DHA ratio (2:1), which might affect the offspring’s growth and adiposity.

We presume that maternal n-3 PUFA deficiency affects the hippocampal expression of neurogenesis genes due to altered mammary lipid composition in the dam. The disrupted expression of neurogenesis genes and energy transporter (GLUTs) in the offspring’s brain could affect functionalities as both are required for neuronal firing during learning.

## Abbreviations

n-3 PUFA: n-3 (omega-3) polyunsaturated fatty acid
LA: Linoleic acid
ALA: Alpha-linolenic acid
LCPUFA: Long-chain polyunsaturated fatty acid
ARA: Arachidonic acid
DHA: Docosahexaenoic acid
EPA: Eicosapentaenoic acid
AIN 93: American institute of nutrition 93
gD: Gestational day
FAME: Methyl esters of fatty acids
gDNA: Genomic deoxyribonucleic acid
cDNA: Complementary deoxyribonucleic acid
mRNA: Messenger ribonucleic acid
RT-qPCR: Quantitative reverse transcription polymerase chain reaction
Ct: Threshold cycle
kcal: kilocalorie

## Author declarations

Competing interests - The authors declare no competing interests.

Ethics approval - Compliance with ethical standards

Consent to participate - Not applicable.

Consent for publication - Not applicable.

Availability of data and material-Data Transparency

Code Availability - Not applicable.

## Acknowledgements

The study was sponsored by the fund received from the Department of Biotechnology, Government of India (BT/PR6946/MED), and ICMR-National Institute of Nutrition (grant number 17-BS05).

## Contribution statement

VS conducted the animal trial, sample and data collection, performed significant laboratory experiments, data collection; SV was involved in mRNA expression by RT-qPCR; SRK was involved in diet preparation and fatty acid composition; AI provided inputs on study design, reviewed, commented on the manuscript; AKDR provided critical comments and review of the manuscript. SB conceptualized the study, designed, analyzed, drafted, interpreted and finalized the manuscript.

## Notes

### Competing Interest Statement

The authors have declared no competing interest.

### Summary of Updates

Author names are updated

