## Supplementary material for "Dietary omega-3 fatty acid deficiency from pre-pregnancy to lactation affects expression of genes involved in neurogenesis of the offspring": Sup Table 1

Supplementary Table 1: Genes studied in mouse neurogenesis by RT^2^ profiler qRT-PCR array

| **GenBank ID** | **Gene Symbol** | **Gene name** |
| --- | --- | --- |
| NM_009599 | ACHE | Acetylcholinesterase |
| NM_001008533 | ADORA1 | Adenosine A1 receptor |
| NM_009630 | ADORA2A/A2AAR | Adenosine A2a receptor |
| NM_007439 | ALK | Anaplastic lymphoma kinase |
| NM_009685 | APBB1 | Amyloid beta(A4) precursor protein-binding, family B, member 1 |
| NM_009696 | APOE | Apolipoprotein E |
| NM_007471 | APP | Amyloid beta(A4) precursor protein |
| NM_009711 | ARTN | Artemin |
| NM_008553 | ASCL1 | Achaete-scute complex homolog1(Drosophila) |
| NM_009741 | BCL2 | B-cell leukemia/lymphoma2 |
| NM_007540 | BDNF | Brain derived neurotrophic factor |
| NM_007553 | BMP2 | Bone morphogenetic protein 2 |
| NM_007554 | BMP4 | Bone morphogenetic protein 4 |
| NM_007559 | BMP8B | Bone morphogenetic protein 8b |
| NM_009871 | CDK5R1 | Cyclin-dependent kinase 5, regulatory subunit 1 (p35) |
| NM_145990 | CDK5RAP2 | CDK5 regulatory subunit associated protein 2 |
| NM_203491 | CHRM2 | Cholinergic receptor muscarinic 2, cardiac |
| NM_133828 | CREB1 | CAMP responsive element binding protein 1 |
| NM_008176 | CXCL1 | Chemokine (CXC motif) ligand 1 |
| NM_010025 | DCX | Doublecortin |
| NM_007864 | DLG4 | Discs large homlog 4 (Droshophila) |
| NM_007865 | DLL1 | Delta-like 1 (Drosophila) |
| NM_010077 | DRD2/D2R | Dopamine receptor D2 |
| NM_007889 | DVL3 | Dishevelled Segment Polarity Protein 3 (Drosophila) |
| NM_010110 | EFNB1 | Ephrin B1 |
| NM_010113 | EGF | Epidermal growth factor |
| NM_177821 | EP300 | E1A binding protein, p300 |
| NM_001003817 | ERBB2 | erb-b2 glioblastoma derived oncogene homolog (avian) |
| NM_008006 | FGF2 | Fibroblast growth factor 2 |
| NM_010227 | FLNA | Filamin alpha |
| NM_010275 | GDNF | Glial cell line derived neurotrophic factor |
| NM_008155 | GPI1 | Glucose phosphate isomerase 1 |
| NM_008169 | GRIN1 | Glutamate receptor, ionotropic NMDA1(zeta 1) |
| NM_207225 | HDAC4 | Histone deacetylase 4 |
| NM_008235 | HES1 | Hairy and enhancer of split 1(drosophila) |
| NM_010423 | HEY1 | Hairy/enhancer-of-split related with YRPW motif 1 |
| NM_013904 | HEY2 | Hairy/enhancer-of-split related with YRPW motif 2 |
| NM_013905 | HEYL | Hairy/enhancer-of-split related with YRPW motif –like |
| NM_010556 | IL3 | Interleukin 3 |
| NM_010784 | MDK | Midkine |
| NM_025282 | MEF2C | Myocyte enhancer factor 2c |
| NM_001081049 | MLL1 | Myeloid/lymphoid or mixed-lineage leukemia 1 |
| NM_001039934 | MTAP2 | Microtubule-associated protein 2 |
| NM_010882 | NDN | Necdin |
| NM_010883 | NDP | Norrie disease, pseudoglioma (human) |
| NM_010894 | NEUROD1 | Neurogenic differentiation 1 |
| NM_010896 | NEUROG1 | Neurogenin 1 |
| NM_009718 | NEUROG2 | Neurogenin 2 |
| NM_010897 | NF1 | Neurofibromatosis 1 |
| NM_008711 | NOG | Noggin |
| NM_008714 | NOTCH1 | Notch gene homolog 1 (Drosophila) |
| NM_010928 | NOTCH2 | Notch gene homolog 2 (Drosophila) |
| NM_013708 | NR2E3 | Nuclear receptor sub family 2, group E, member 3 |
| NM_176930 | NRCAM | Neuron-glia-CAM-related cell adhesion molecule |
| NM_178591 | NRG1 | Neuregulin 1 |
| NM_008737 | NRP1 | Neuropilin 1 |
| NM_010939 | NRP2 | Neuropilin 2 |
| NM_008742 | NTF3 | Neurotrophin 3 |
| NM_008744 | NTN1 | Netrin 1 |
| NM_011855 | ODZ1 | Odd Oz/ten-m homolog 1 (Drosophila) |
| NM_016967 | OLIG2 | Oligodendrocyte transcription factor 2 |
| NM_013625 | PAFAH1B1 | Platelet-activating factor acetyl hydrolase, isoform 1b, subunit 1 |
| NM_033620 | PARD3 | Par-3 (partitioning defective 3) homolog (C. elegans) |
| NM_008781 | PAX3 | Paired box gene 3 |
| NM_008782 | PAX5 | Paired box gene 5 |
| NM_013627 | PAX6 | Paired box gene 6 |
| NM_008900 | POU3F3 | POU Class 3 Homeobox 3 |
| NM_011143 | POU4F1 | POU Class 4 Homeobox 1 |
| NM_008973 | PTN | Pleiotrophin |
| NM_009007 | RAC1 | RAS-related C3 Botulinum substrate 1 |
| NM_019413 | ROBO1 | Roundabout homolog 1 (Drosophila) |
| NM_194053 | RTN4 | Reticulon 4 |
| NM_011313 | S100A6 | S100 calcium binding protein A6 (calcyclin) |
| NM_009115 | S100B | S100 protein Beta polypeptide neural |
| NM_009170 | SHH | Sonic hedgehog |
| NM_178804 | SLIT2 | Slit homolog 2 (Drosophila) |
| NM_011434 | SOD1 | Superoxide dismutase 1, soluble |
| NM_011443 | SOX2 | SRY-box containing gene 2 |
| NM_009237 | SOX3 | SRY-box containing gene 3 |
| NM_011486 | STAT3 | Signal transducer and activator of transcription 3 |
| NM_011577 | TGFB1 | Transforming growth factor, beta 1 |
| NM_009377 | TH | Tyrosine hydroxylase |
| NM_022312 | TNR | Tenascin R |
| NM_009505 | VEGFA | Vascular endothelial growth factor A |
| NM_007393 | ACTB | Actin beta |
| NM_009735 | B2M | Beta-2, macroglobulin |
| NM_008084 | GAPDH | Glyceraldehyde-3-phosphate dehydrogenase |
| NM_010368 | GUSB | Glucuronidase beta |
| NM_008302 | HSP90AB1 | Heat shock protein 90 alpha (cytosolic class B member 1) |
